# Timing and localization of myasthenia gravis-related gene expression

**DOI:** 10.1101/2021.01.05.425366

**Authors:** Dana L.E. Vergoossen, Arlin Keo, Ahmed Mahfouz, Maartje G. Huijbers

## Abstract

Myasthenia gravis (MG) is an acquired autoimmune disorder caused by autoantibodies binding acetylcholine receptors (AChR), muscle-specific kinase (MuSK), agrin or low-density lipoprotein receptor-related protein 4 (Lrp4). These autoantibodies inhibit neuromuscular transmission by blocking the function of these proteins, and thereby cause fluctuating skeletal muscle weakness. Several reports suggest that these autoantibodies might also affect the central nervous system (CNS) in MG patients. A comprehensive overview of the timing and localization of the expression of MG-related antigens in other organs is currently lacking. To investigate the spatio-temporal expression of MG-related genes outside skeletal muscle, we used *in silico* tools to assess public expression databases. Acetylcholine esterase, nicotinic AChR α1 subunit, agrin, collagen Q, Dok7, Lrp4, MuSK and rapsyn were included as MG-related genes because of their well-known involvement in either congenital or autoimmune MG. We investigated expression of MG-related genes in 1) all human tissues using GTEx data, 2) specific brain regions, 3) neurodevelopmental stages, and 4) cell types using datasets from the Allen Institute for Brain Sciences. MG-related genes show heterogenous spatio-temporal expression patterns in the human body as well as in the CNS. For each of these genes several (new) tissues, brain areas and cortical cell types with (relatively) high expression were identified suggesting a potential role for these genes outside skeletal muscle. The possible presence of MG-related antigens outside skeletal muscle suggests that autoimmune MG, congenital MG or treatments targeting the same proteins may affect MG-related protein function in other organs.

## Introduction

Myasthenia gravis (MG) is an antibody-mediated autoimmune disorder hallmarked by fatigable skeletal muscle weakness. This muscle weakness results from autoantibodies targeting essential proteins at the neuromuscular junction (NMJ). Till date, four antigens have been described: acetylcholine receptors (AChR), muscle-specific kinase (MuSK), low-density lipoprotein receptor-related protein 4 (Lrp4) and agrin (1). These proteins converge on a single pathway essential for establishing and maintaining NMJs and facilitating neuromuscular transmission (2). Consequently, binding of autoantibodies inhibits the function of these proteins, resulting in failure of neuromuscular transmission and subsequent muscle fatigue and paralysis.

Even though most knowledge on these proteins relates to their function in the NMJ, they are also expressed in tissues other than skeletal muscle, like retina, lung and brain (2–5). Insight in the localization of expression of these genes/proteins and their roles in other organs is important because: 1) Other tissues may also be affected by the autoantibodies in MG patients and 2) new therapeutic strategies targeting these MG-related genes/proteins are immerging and therefore knowledge on their localization may identify potential off-target effects. A comprehensive overview of the expression of MG-related genes in different human tissues however is lacking.

Evidence of other organs being affected by MG autoantibodies is mostly focussed on the central nervous system (CNS). The non-motor symptoms include pain, cognitive dysfunction, fatigue and sleep disturbances (6–10). Although AChR MG serum antibodies were reported not to bind neuronal AChR (11) and immunostaining with MuSK autoantibodies seems challenging on brain sections (personal observation), passive transfer of patient-derived MuSK and AChR antibodies resulted in behavioural deficits and EEG abnormalities in mice (12, 13). Moreover, AChR autoantibodies have been detected in cerebrospinal fluid (14–16). Mutations in NMJ genes may furthermore result in congenital myasthenic syndrome (CMS) and sometimes give CNS abnormalities (17). Although CNS defects are not at the foreground of clinical symptoms in MG patients, these observations suggest that autoantibodies may have detrimental effects in the CNS when they are able to cross the blood-brain barrier.

If indeed MG-related autoantibodies are able to affect their antigens in other organs several important questions arise: e.g. 1) do these proteins, like in the NMJ, converge on a similar pathway in these organs and 2) which cells are responsible for this expression. We therefore investigate the spatio-temporal expression patterns of MG-related genes in healthy human tissues, with a focus on the CNS, using a range of publicly-available expression databases. We selected the four NMJ genes which encode known antigens for autoantibodies in MG (*AGRN, CHRNA1, LRP4, MUSK*) and four other NMJ genes involved in maintaining NMJ neurotransmission and where mutations can cause CMS (acetylcholine esterase (*ACHE*), collagen Q (*COLQ*), downstream of kinase-7 (*DOK7*), and rapsyn (*RAPSN*)) (Table 1) (17). We will refer to these eight genes and their gene products as MG-related genes.

**Table 1.** The MG-related genes, their function at the NMJ and association with autoimmune MG or CMS (1, 2, 17, 47).

| **Gene symbol** | **Encoded Protein** | **Function at NMJ** | **Role in CMS or autoimmune MG** |
| --- | --- | --- | --- |
| *ACHE* | Acetylcholine esterase | Breaks down the neurotransmitter acetylcholine (ACh), thereby halting neuromuscular transmission and muscle contraction. | 12.53% of CMS patients have a AChE deficiency. AChE is the main target for symptomatic treatment of autoimmune MG patients. |
| *AGRN* | Agrin | Trophic signalling molecule released by the presynaptic motor nerve terminal to induce and maintain post-synaptic differentiation. Agrin lingers in the basal lamina, can bind Lrp4 and thereby activates MuSK and AChR clustering. | Mutations in *AGRN* are a rare cause of CMS (0.58%). Autoantibodies to agrin are thought to cause autoimmune MG in a small subset of patients. |
| *CHRNA1* | Nicotinic acetylcholine receptor $\alpha$ subunit | Muscle-specific subunit of the AChR required for neuromuscular transmission and location for ACh binding. Upon binding of ACh to the AChR, its ion channel opens which depolarizes the muscle end plate, and may induce an action potential and muscle contraction. | ~50% of CMS patients have an AChR deficiency caused by mutations in one of the AChR subunits leading to kinetic defects and myasthenic symptoms. ~80% of autoimmune MG patients have autoantibodies to the AChR. The $\alpha$ subunit contains the main immunogenic region. |
| *COLQ* | Collagen Q | Collagenic subunit anchoring AChE to the basal lamina and thereby responsible for ACh breakdown. Also interacts with MuSK. | The pathomechanism of AChE and MuSK CMS as well as autoimmune MG are known to affect ColQ function. |
| *DOK7* | Downstream of kinase 7 | Cytoplasmic adaptor of MuSK required for regulating the kinase activity of MuSK, subsequent AChR clustering and NMJ formation. | 9.75% of CMS patients have a mutation in *DOK7*. *DOK7* gene therapy rescued the phenotype of a Dok7 CMS mouse model. |
| *LRP4* | low-density lipoprotein receptor-related protein 4 | The agrin receptor that directly interacts with MuSK and further activates AChR clustering and postsynaptic differentiation. Lrp4 is also critical for presynaptic differentiation. | 0.56% of CMS patients have mutations in *LRP4*. 1-2% of MG patients have Lrp4 autoantibodies. MuSK autoantibodies inhibit MuSK-Lrp4 interaction, obstructing normal trophic signalling at the NMJ and induce myasthenia. |
| *MUSK* | Muscle-specific kinase | Orchestrates anterograde trophic signalling at the NMJ, resulting in AChR clustering. It is also required for retrograde signalling towards presynaptic differentiation of the motor neuron end plate. | 0.28% of CMS patients have a mutation in *MUSK*. Autoantibodies to MuSK cause autoimmune MG in 5-8% of patients. |
| *RAPSN* | Rapsyn | Cytoplasmic anchor that facilitates AChR clustering. | 14.21% of CMS patients has a rapsyn deficiency. |

## Materials and methods

### Genotype-Tissue expression consortium (GTEx)

The expression of MG-related genes was analysed across 54 non-diseased human tissues of nearly 1000 individuals using RNA sequencing data from the open access Genotype-Tissue expression consortium (GTEx) version 8. For *ACHE* (ENSG00000087085) and *AGRN* (ENSG00000188157), all isoforms were annotated using the Ensembl database. The median gene-level transcripts per million (TPM) for all MG-related genes were downloaded per tissue. Because it is known that *ACHE* and *AGRN* produce different splice variants, of which only one is relevant for NMJs, *ACHE-207* (ENST00000428317.5) and *AGRN-208* (ENST00000620552.4) isoform data were downloaded separately. The anterior cingulate cortex, frontal cortex and cerebellar hemisphere were removed because they were covered in the cortex or cerebellum respectively. EBV-transformed lymphocytes and cultured fibroblasts were removed, because they do not naturally occur in healthy humans. Tissues (rows) and genes (columns) were hierarchically clustered using hclust function in R with Eucledian distance and average linkage. Qualitative expression levels are used as follows: low (Log10 <1), moderate (Log10 1-1.5), high (Log10 >1.5).

### Allen Human Brain Atlas

The Allen Human Brain Atlas (AHBA) includes anatomically-mapped Oligo-dT primed microarray data of 3,702 samples from six neurotypical individuals (5 males and 1 female, mean age 42, range 24–57 years) (18). The expression of the MG-related genes was mapped across regions of the human brain using BrainScope (19). The MG-related genes probes used are marked in Supplemental Table 1. All probes were confirmed to align to their respective MG-related gene using the Ensembl database. *AGRN* probe A_24_P358462 (#) detected the NMJ-specific *AGRN-208* isoform able to induce AChR clustering and clusters separately from the probe used in the other analyses (*) (Supplemental Fig. S3a) (20). For *ACHE*, the available probe did not selectively bind to *ACHE-207*. The expression of MG-related genes was downloaded from http://human.brain-map.org/. Log2-transformed expression values were converted into z-scores normalized per donor. The median z-score of each gene was taken across the set of 22 non-overlapping brain regions. Finally, the median z-score of every MG gene across the six donors was taken. Hierarchical clustering was done as described above. The nature of this data only allowed comparison of z-scores across anatomical brain areas within one gene, which is why the term relative expression is used. Qualitative expression levels are used as follows: low (Z < 0), moderate (Z 0-1), high (Z >1).

### BrainSpan atlas of the developing human brain

The BrainSpan atlas of the developing human brain includes 42 healthy human brains, ranging in age from 8 weeks post-conception to 40 years old, from which a total of 524 anatomically annotated samples were taken (21). Gene expression was determined using RNA sequencing and visualized using BrainScope (19). *ACHE-207* and *AGRN-208* isoforms could not be distinguished in this dataset. MG-related genes *CHRNA1, DOK7, MUSK* and *RAPSN* were not present in BrainScope, since genes with RPKM-value above 1 in less than 20% of all samples were removed. For those genes, we used the brain-map.org portal to plot their expression (Supplemental Fig. 4).

### Single-cell analyses

Single-nucleus RNA sequencing (snRNA-seq) data from multiple human cortical regions and single-cell RNA sequencing data from the whole mouse cortex were downloaded from the Allen Institute (https://portal.brain-map.org/atlases-and-data/rnaseq). For the human dataset, the donors included 4 males and 4 females (age 24-66) without a history of neuropsychiatric or neurological conditions (22). From 4 donors, multiple cortical areas were sampled postmortem and from 4 donors the medial temporal gyrus was removed during neurosurgery. For the mouse dataset, 538 animals were used from multiple transgenic lines to enrich for rare cell types, all on a C57BL/6J background (23).

Sample processing and analysis methods have been described previously (22, 23). Briefly, the SMART-seq method yielded transcriptome profiles for 10,708 glutamatergic neurons, 4,297 GABAergic neurons and 923 non-neuronal cells for the human dataset and 40,276 glutamatergic neurons, 22,573 GABAergic neurons and 1,958 non-neuronal cells for the mouse dataset. The trimmed-mean gene expression (CPM) data per cluster was downloaded for 8 MG-related genes and grouped in the cell type subclasses defined in the sampling strategy. In the mouse dataset, we selected the analogous cell type subclasses that are present in the human dataset (22).

### Code and Data Availability

Scripts to generate all the results presented in this manuscript can be found online at: https://github.com/ahmedmahfouz/MG-analysis.

## Results

### Expression of MG-related genes in muscle and brain regions

Expression of MG-related genes was assessed in human tissues using GTEx data. For *ACHE* and *AGRN*, specific splice variants (*ACHE-207* and *AGRN-208*) are known to be (NMJ) synapse-specific (20, 24). *ACHE-207* and *AGRN-208* were indeed specifically expressed in skeletal muscle and/or brain regions, in contrast to other isoforms (Supplemental Fig. 1). Because of their relevance for the NMJ and association with autoimmune MG, *ACHE-207* and *AGRN-208* were selected for further analysis (Fig. 1). For completeness, the analysis was also performed using gene counts (i.e. including all isoforms) in Supplemental Fig. 2.

**Fig. 1.**
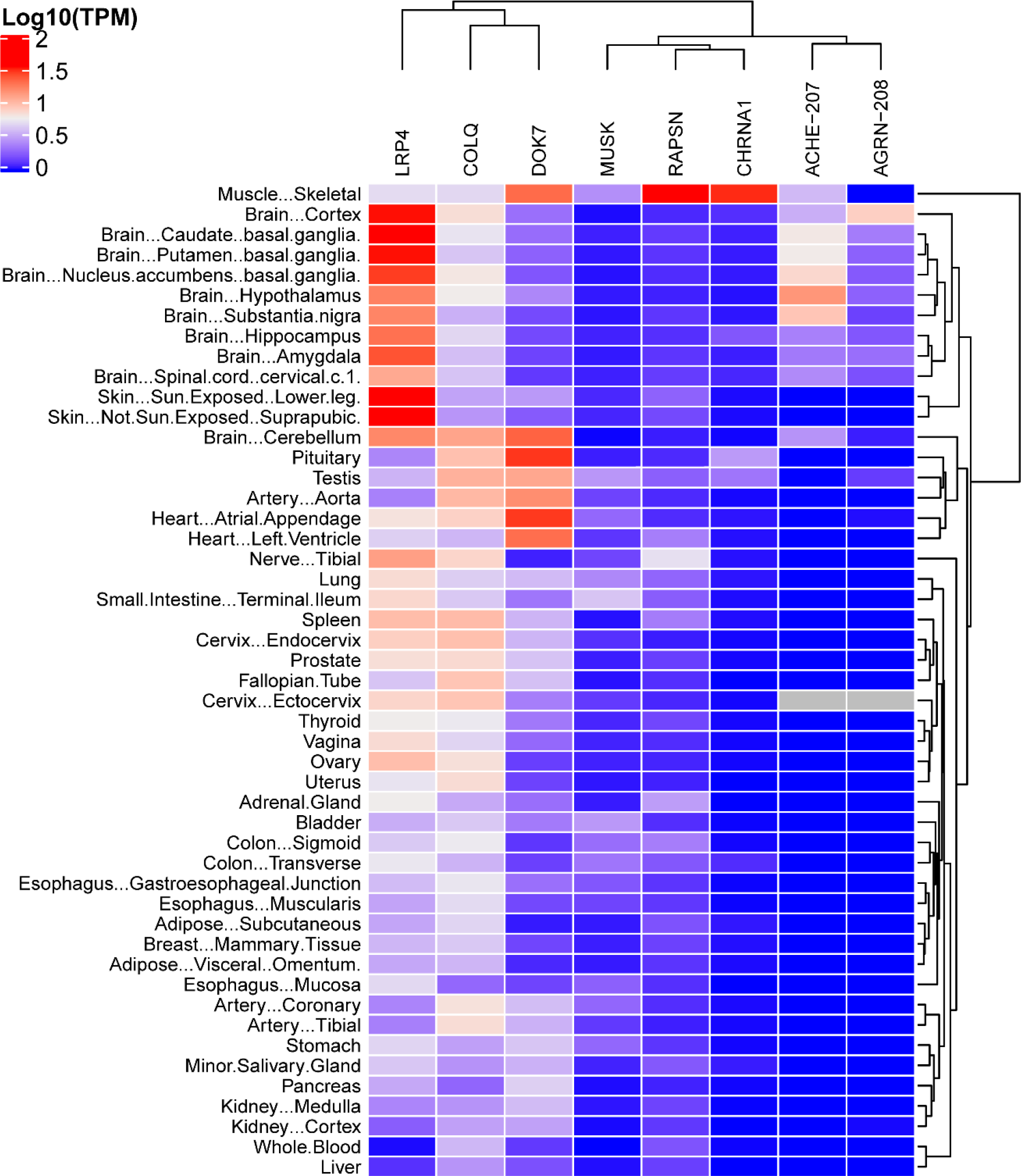
Overview of tissue-specific expression of MG-related genes. reveals most co-expression in skeletal muscle, the CNS and parts of the male and female reproductive system. The heatmap shows the median expression of each MG-related gene (column) in TPM across human tissues (rows) using data from GTEx V8. Average expression is reported as Log10(TPM+1).

Hierarchical clustering revealed that skeletal muscle segregates from all other tissues based on expression of MG-related genes, supporting the unique combined role of these genes in the NMJ (Fig. 1). Agrin is secreted by motor neurons, explaining the absence of *AGRN-208* expression in skeletal muscle. The absence of *AGRN-208* in the tibial nerve may be explained by localization of mRNA to subcellular compartments in long axons (25). Such transcripts are missed when sampling the nerve nucleus.

Cerebral brain areas clustered together close to skeletal muscle with expression of nearly all MG-related genes, confirming expression these genes in healthy adult human brain. All MG-related genes could be detected in areas such as hippocampus and basal ganglia, although the amount of expression differed compared to skeletal muscle. The cerebellum clustered separately from the cerebrum with moderate expression of *DOK7, LRP4* and *COLQ*. Transcriptomic separation of cerebellum and cerebrum is observed for many other genes (19). Outside the brain, almost all MG-related genes are found in the testis and the ectocervix. Other components of the male and female reproductive systems cluster together with moderate expression of *LRP4* and *COLQ* and low expression of *DOK7*, *MUSK* and/or *RAPSN*. Varying expression for subsets of MG-related genes could be detected in the remaining tissues.

We observed three clusters of the MG-related genes (Fig. 1). *LRP4, COLQ* and *DOK7* have relatively ubiquitous expression across all tissues. Notably, *LRP4* was most expressed in the brain and the skin, *COLQ* in the cerebellum, pituitary gland, testis and heart, and *DOK7* in the heart and pituitary gland. In contrast, *MUSK, RAPSN* and *CHRNA1* showed low expression in a more limited subset of tissues. *MUSK* expression is highest in the small intestine, bladder and testis. *RAPSN* and *CHRNA1* expression was largely restricted to skeletal muscle, with some additional expression in the tibial nerve, and testis and pituitary gland respectively. Finally, the *ACHE-207* and *AGRN-208* isoforms clustered together based on their expression in the brain and ectocervix. Taken together, the expression of MG-related genes is prominent in skeletal muscle and the brain, but individual genes are also expressed in other tissues of the human body.

### NMJ genes do not share anatomical expression patterns in the brain

Compared to the ten brain regions in the GTEx database, the AHBA provides a high-resolution map of relative gene expression across the adult human brain. Strong correlation between spatial expression patterns would become apparent when genes cluster in the same region of the t-SNE plot (Fig. 2a). However, MG-related genes are scattered across the t-SNE plot, suggesting they do not share spatial expression patterns across the brain.

**Fig. 2.**
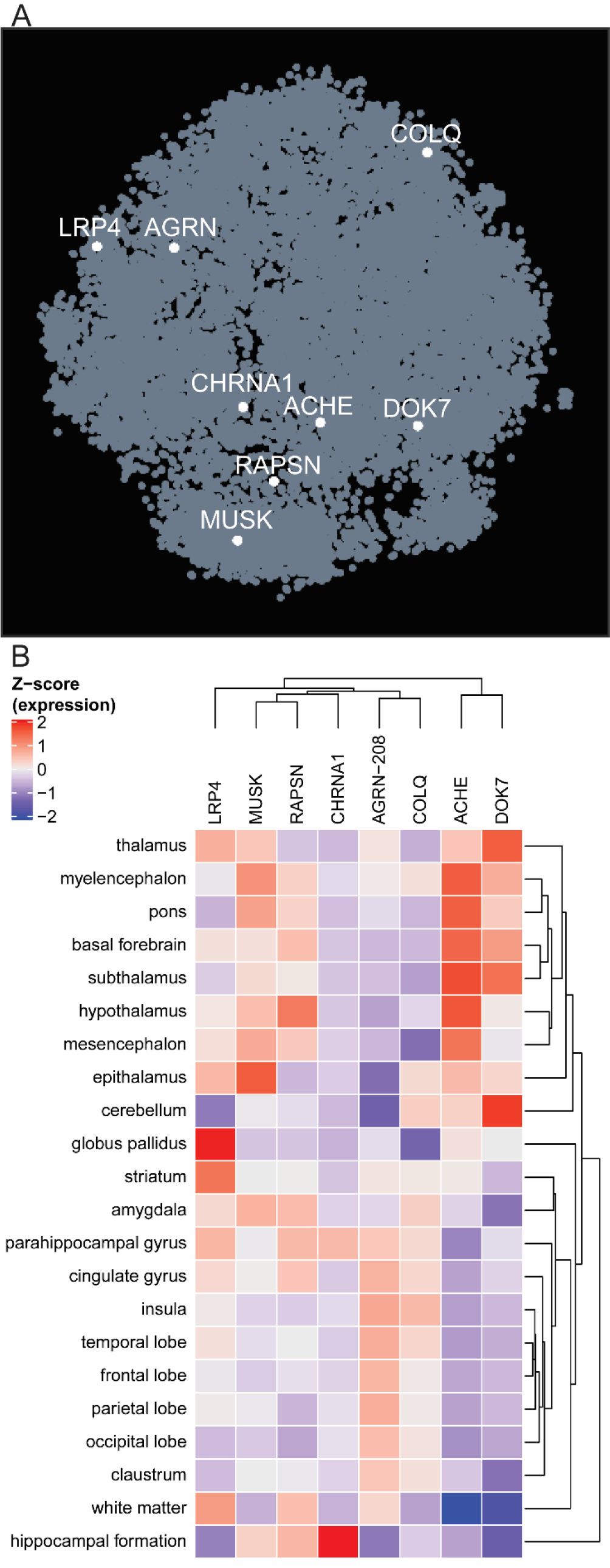
Spatial expression of MG-related genes in the adult human brain. Co-expression analysis of MG-related genes, based on their distribution in a t-SNE embedding, reveals that they do not share spatial expression (a). The t-SNE from BrainScope shows an embedding of all genes (dots) based on their expression pattern across the whole brain. b A heatmap showing the average expression of MG genes across distinct (nonoverlapping) brain regions. Expression values represent the average z-score normalized expression per gene (i.e. relative expression across brain regions).

For hierarchical clustering analysis of distinct brain regions, the NMJ-specific isoform of agrin (*AGRN-208*) was included (Fig. 2b) while a heatmap including all *AGRN* isoforms can be found in Supplemental Fig. S3b. The hippocampus clustered separately from all other brain regions with the unique pattern of high relative expression of *CHRNA1, RAPSN* and *MUSK* (Fig. 2b). The remaining anatomical brain regions separated in two large clusters enriched in the developmentally early and late regions respectively. The early developmental cluster with hindbrain, midbrain and diencephalon was characterized by high relative expression of *DOK7* and *ACHE* and predominant low relative expression of *COLQ* and *AGRN-208*. In contrast, the late developmental cluster, with the basal ganglia, limbic system and cortical regions, was characterized by moderate relative expression of *COLQ* and *AGRN-208* and low relative expression of *DOK7* and *ACHE*. Furthermore, expression is highest in a unique structure for most genes. *DOK7* in cerebellum, *CHRNA1* in hippocampus, *MUSK* in epithalamus, *LRP4* in globus pallidus and *RAPSN* in the hypothalamus. Taken together, this suggests that MG-related genes are likely not active in similar pathways in the CNS as they do not share spatial expression patterns. However, subsets of these genes may be involved in signalling in similar structures.

### NMJ genes do not co-segregate in developmental time in the human brain

To investigate whether expression of these genes follows a temporal pattern in brain development, we used the BrainSpan atlas visualized in the BrainScope browser (19, 21). The expression of *ACHE, AGRN, COLQ* and *LRP4* did not follow the same pattern across human brain development (Fig. 3). *AGRN* was particularly expressed until early childhood while *LRP4* was expressed from the 3^rd^ prenatal stage into adulthood. *ACHE* expression was predominantly restricted to the cerebellum, thalamus, amygdala and striatum from birth until adulthood. *COLQ* is highest across the entire brain in adult life, although various brain regions also show high expression in the last prenatal stage. *CHRNA1, DOK7, MUSK* and *RAPSN* could not be studied with BrainScope, but the brain-map.org portal confirms the low and unique expression patterns of these genes (Supplemental Fig. 4). In sum, spatio-temporal expression of MG-related genes is heterogenous in the human brain.

**Fig. 3.**
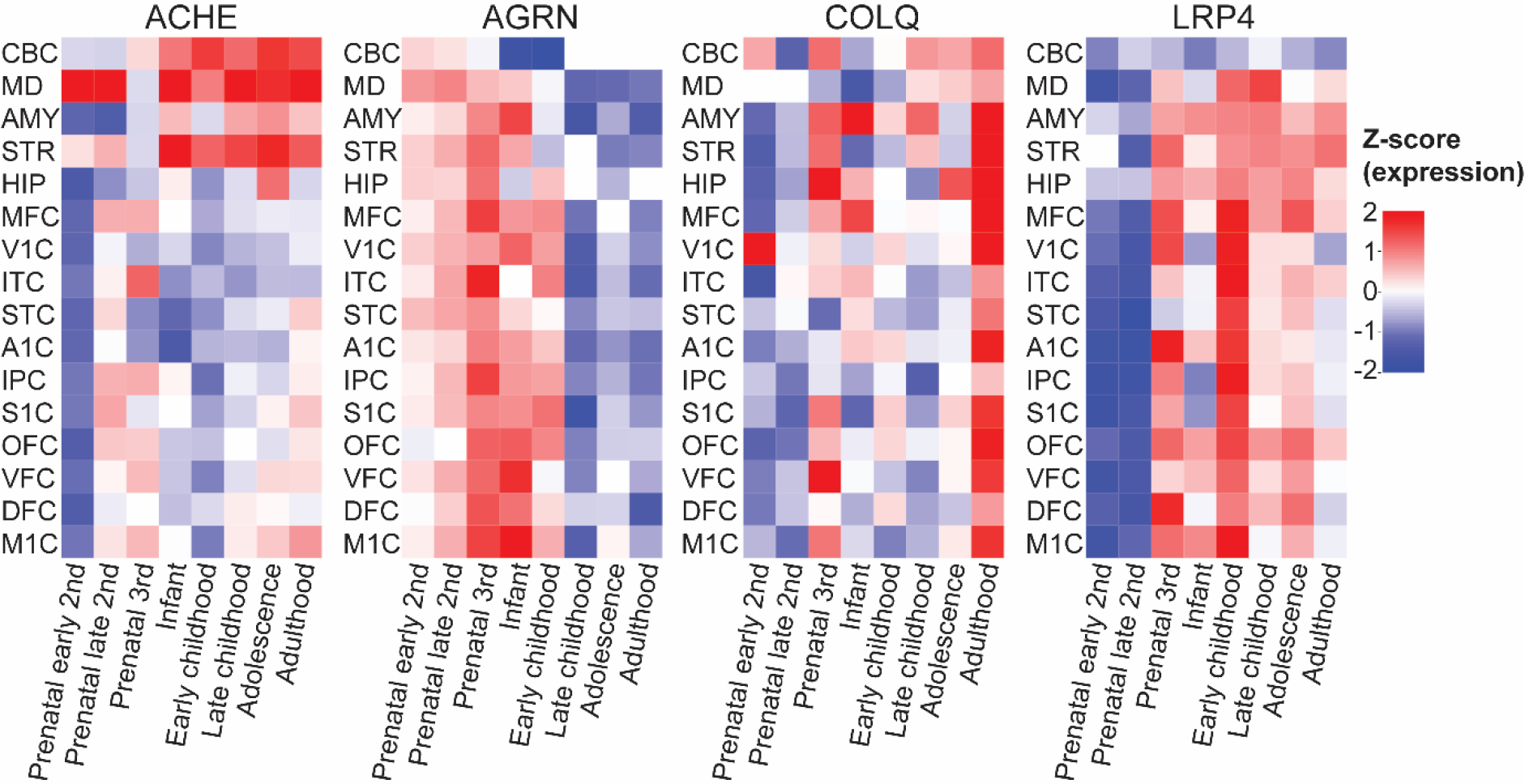
Overview of the expression patterns of four MG-related genes throughout human brain development. *ACHE, AGRN, COLQ* and *LRP4* are expressed during different phases of human brain development. CBC = cerebellar cortex, MD = mediodorsal nucleus of the thalamus, AMY = amygdaloid complex, STR = striatum, HIP = hippocampal formation, MFC = medial prefrontal cortex, V1C = primary visual cortex, ITC = inferolateral temporal cortex, STC = superior temporal cortex, A1C = primary auditory cortex, IPC = interior parietal cortex, S1C = primary somatosensory cortex, OFC = orbital frontal cortex, VFC = ventrolateral prefrontal cortex, DFC = dorsolateral prefrontal cortex, M1C = primary motor cortex.

### MG-related genes do not co-express in the same cell types in the cortex

To understand which cell types are responsible for the regional expression of MG-related genes in the CNS, we explored a snRNAseq dataset (22). The expression of *CHRNA1* and *RAPSN* could not be detected in cortical cell types (Fig. 4a). This is in line with the very low to absent expression in the bulk RNAseq datasets of the cortical regions (Fig. 1 and Supplemental Fig. 4). Interestingly, *MUSK* and *LRP4* were uniquely expressed in nonneuronal cell types, with *MUSK* solely in vascular leptomeningeal cells (VLMC) and oligodendrocytes and *LRP4* predominantly in astrocytes and oligodendrocytes progenitor cells (Fig. 4a). *ACHE*, *AGRN*, *COLQ* and *DOK7* were predominantly expressed in glutamatergic and/or GABAergic neurons; although, *AGRN* was also expressed in pericytes. Overall, a variety of cell types seems to be responsible for expression of MG-related genes in the cortex. As many preclinical studies related to MG are done in rodents, it is relevant to know how the expression of MG-related genes compares between humans and rodents. In mouse cortical regions similar cell types expressed *MUSK*, *LRP4*, *CHRNA1* and *RAPSN*, suggesting adequate translatability of results (Fig. 4b). In contrast, *ACHE* and *AGRN* expression was ubiquitously high in mouse neuronal cell types, but much more restricted to GABAergic or glutamatergic cell types respectively in the human cortex. For *COLQ*, the expression is ubiquitously high in human neuronal cell types, but very much restricted to a single glutamatergic subclass in the mouse. For these genes, translatability of functional studies to humans may thus be limited.

**Fig. 4.**
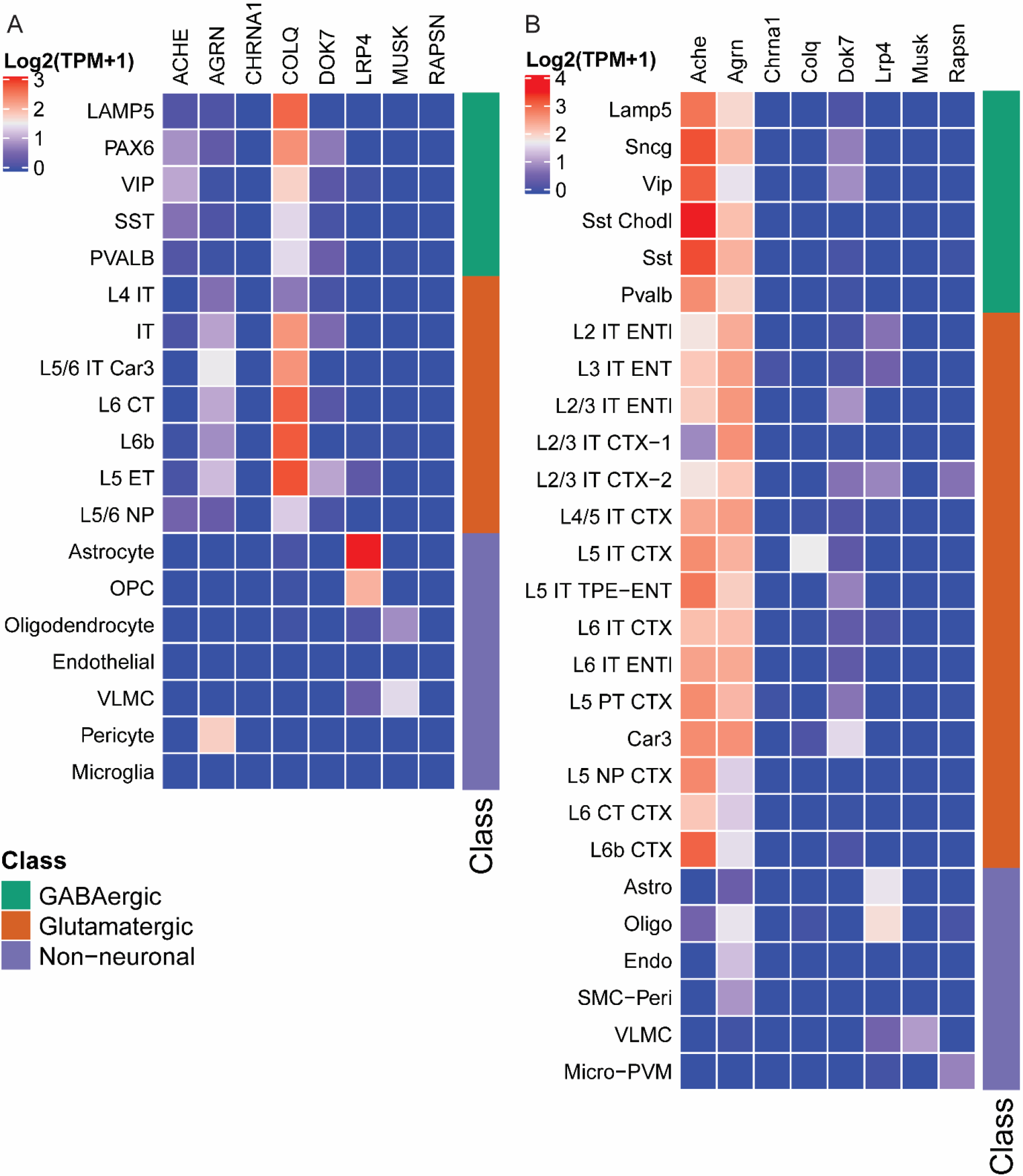
Cortical cell type expression of MG-related genes. Heatmap of adult human cortical cell types (a). Heatmap of adult mouse cortical cell types (b). VLMC = ventricular leptomeningeal cells, OPC = oligodendrocyte progenitor cells.

## Discussion

This study confirms that many of the MG-related genes are widely expressed outside skeletal muscle, with the exact pattern of involved organs varying per gene. Within the CNS, these genes furthermore show heterogenous spatial, temporal and cellular expression patterns. This suggests that these genes do not converge on a single pathway in the brain or other human tissues. It furthermore provides valuable insights for new research questions and hypotheses, explaining possible non-motor symptoms in myasthenic syndromes or off-target effects of MG-related targeted treatments.

Validity of these expression data is supported by studies confirming the presence of these MG-related proteins outside skeletal muscle. Congruent with the observed expression in testis, agrin, MuSK and rapsyn are present in human sperm, (26, 27). Dok7 protein was detected in heart and brain, and MuSK in bladder, heart, lung and liver (28, 29). Agrin is present in glutamatergic neurons and increasingly surrounding brain microvasculature during development of the blood-brain barrier, validating the expression in pericytes and during prenatal development (30, 31). Lrp4 is indeed abundantly present in the brain, most prominently in astrocytes (3, 32). Not all observations however match with previous studies. Dok7 was not detected in liver and spleen before (28, 29) and MuSK protein was detected in spleen, but is not present in our dataset. MuSK and agrin were found in glutamatergic neurons in the mouse cortex, while *MUSK* is only expressed in VLMCs and *AGRN* also in GABAergic neurons in the cell-type analysis (33). It is well established that RNA levels do not always translate to protein; however, discrepancies may also be explained by sensitivity and type of detection method, timing or location of sampling, or differences between rodents and humans. The Allen Institute databases furthermore cover a limited number of individuals, making it difficult to translate these observations to a larger heterogenous population. Cell type-specific expression is currently only available for the cortex, covering a fraction of the cellular complexity in the CNS. More single cell data is anticipated as this field is rapidly expanding. Keeping these limitations in mind, our gene expression analysis method provides biological rationale to further investigate a possible role for candidate genes in newly identified target tissues. This is particularly important for MG-related genes like *Musk, Agrin, Dok7* and *Lrp4*, for which the role in other organs may be overlooked, because null mice die at birth of impaired breathing due to dysfunctional NMJs (28, 34–36).

Surprisingly, many of the MG-related genes were found in parts of the male and female reproductive system. Their role there remains to be uncovered. However, fertility issues have not been described in MG patients nor in mouse models lacking these genes. Interestingly, quite a number of MG-related genes have prominent expression in non-neuronal cell types in the cortex, suggesting a role of these genes outside a synaptic context. Expression of *MUSK* in VLMCs has not been described before and provides an important clue for further research into the role of MuSK in the brain. Till date, for a limited number of MG-related genes the function was studied in the CNS. Reduced agrin, Lrp4 or MuSK levels impaired cortical synaptogenesis and/or hippocampal functioning in mice (29, 32, 33, 37–39). This fits the observation that all three genes are expressed in the hippocampus, although the prominent effects of reducing MuSK or agrin were surprising considering the relatively low expression of *MUSK* and *AGRN-208*. Whether these three genes converge on the same pathway in the hippocampus is uncertain. Agrin was shown to induce MuSK signalling through Lrp4 in hippocampal astrocytes (39). However, the role of agrin and Lrp4 in hippocampal neurogenesis seems to be mediated by Ror2, instead of MuSK, supporting that subsets of MG-related genes may also work together with other proteins (38). Taken together, MG-related genes may participate in similar pathways under certain condition in the hippocampus, but little is known about their role in the rest of the brain. The spatial, temporal and cellular resolution of our analysis can guide future studies to specific anatomical regions, developmental phase and cell type in the brain.

The expression of MG-related genes outside skeletal muscle suggests that other organs may be at risk for impairment by autoantibodies. In autoimmune MG and CMS patients skeletal muscle weakness is clearly at the foreground of symptoms. The presence of these genes in the CNS suggests that the reported CNS-related symptoms in subsets of MG patients may be due to autoantibodies binding their target if they cross the blood-brain barrier (6–10). In CMS patients a mutational bias may occur as observed genetic defects in these genes are likely to result in a sufficiently mild phenotype to allow development. Absence of other symptoms may be explained by MG-related gene products 1) having low levels of expression, 2) are inaccessible to autoantibodies, 3) do not fulfil an essential function, 4) have alternative splice variants or 5) are differentially post-translationally modified masking relevant epitopes. Furthermore, observed non-motor symptoms or comorbidities may also be due to underlying immune dysfunction, thymoma’s or treatment side-effects. Future studies are needed to investigate whether CNS-related or other non-motor symptoms in MG patients are due to autoantibody binding or gene dysfunction not yet recognized.

MG-related genes or proteins are interesting targets for treatment of neuromuscular diseases through strengthening NMJs and improving or maintaining muscle function (40). *DOK7* gene therapy, agonistic MuSK antibodies and agrin biologicals have proven beneficial in several mouse models for ALS, Dok7 CMS, Emery Dreifuss muscular dystrophy, spinal muscular atrophy or sarcopenia (41–46). Off-target effects in other organs have not been reported, but also not specifically investigated. Our data furthermore identified some differences between mouse and human MG-related gene expression which emphasizes where caution is needed for accurate interpretation and translatability of studies using mouse models. For further clinical development, our data provides clear guidance as to which organs may be at risk for off-target effects. Since many of the MG-related genes are expressed in the brain, a treatment strategy that does not cross the blood-brain barrier is recommended.

This hypothesis-free approach to study timing and localization of MG-related gene expression suggests wide-spread expression of these genes in other organs. These insights can guide future studies to uncover their role outside skeletal muscle and guide pre-clinical development of related novel therapeutics.

## Acknowledgements

Thanks to Mink Schinkelshoek for our fruitful joint discussions on the methodology and scientific relevance and Prof. Jan Verschuuren and Prof. Silvère van der Maarel for their support and advice on the manuscript.

