## Supplemental Material for "Timing and localization of myasthenia gravis-related gene expression"

**Supplemental Table 1.** genes and probes used for MG-related genes from the AHBA. * used for analysis, unless otherwise specified. # detects the AGRN-208 isoform able to induce AChR clustering at the NMJ.

| **Gene names** | **Allen Brain Atlas probes** |
|---|---|
| *ACHE* | A_24_P60845* |
| *AGRN* | A_23_P343411*<br>A_24_P24819<br>A_24_P358462#<br>CUST_72_PI416408490 |
| *CHRNA1* | A_23_P90888*<br>CUST_14430_PI416261804 |
| *COLQ* | A_23_P212126*<br>A_24_P26160 |
| *DOK7* | A_23_P39885*<br>CUST_10065_PI416261804 |
| *LRP4* | A_24_P403561*<br>CUST_9952_PI416261804 |
| *MUSK* | A_23_P71649*<br>A_24_P129536 |
| *RAPSN* | A_23_P86801<br>CUST_899_PI416261804* |

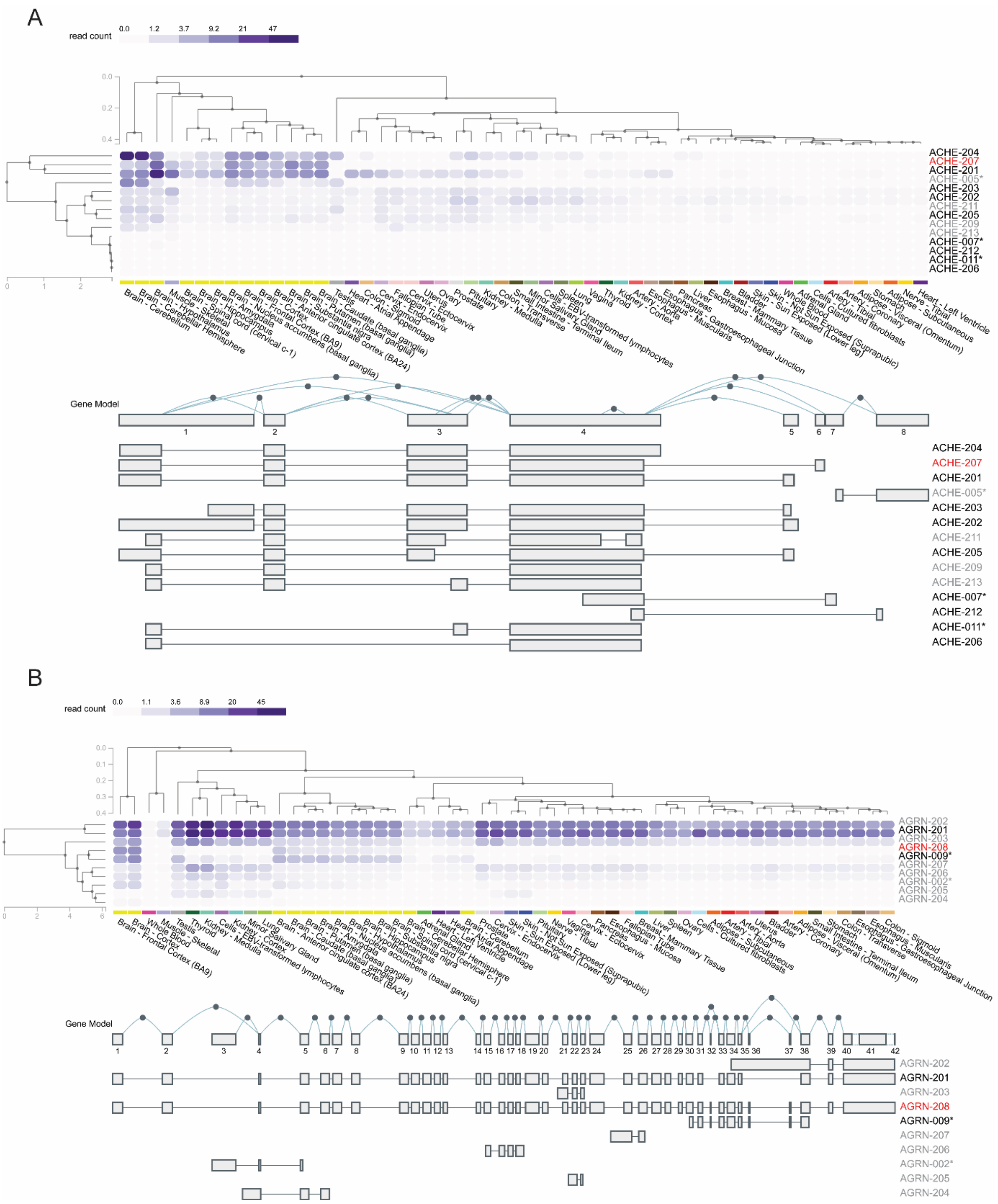


**Supplemental Fig. 1.** Expression of *ACHE* isoforms (A) and *AGRN* isoforms (B) extracted from the GTEx database. Expression of synapses-specific *ACHE-207* isoform (red) is limited to brain areas and skeletal muscle. Expression of NMJ-specific *AGRN-208* isoform (red) is limited to brain areas. Grey isoforms are non-protein coding according to Ensembl (GRCh38). * isoforms identified in earlier assemblies of the human genome.

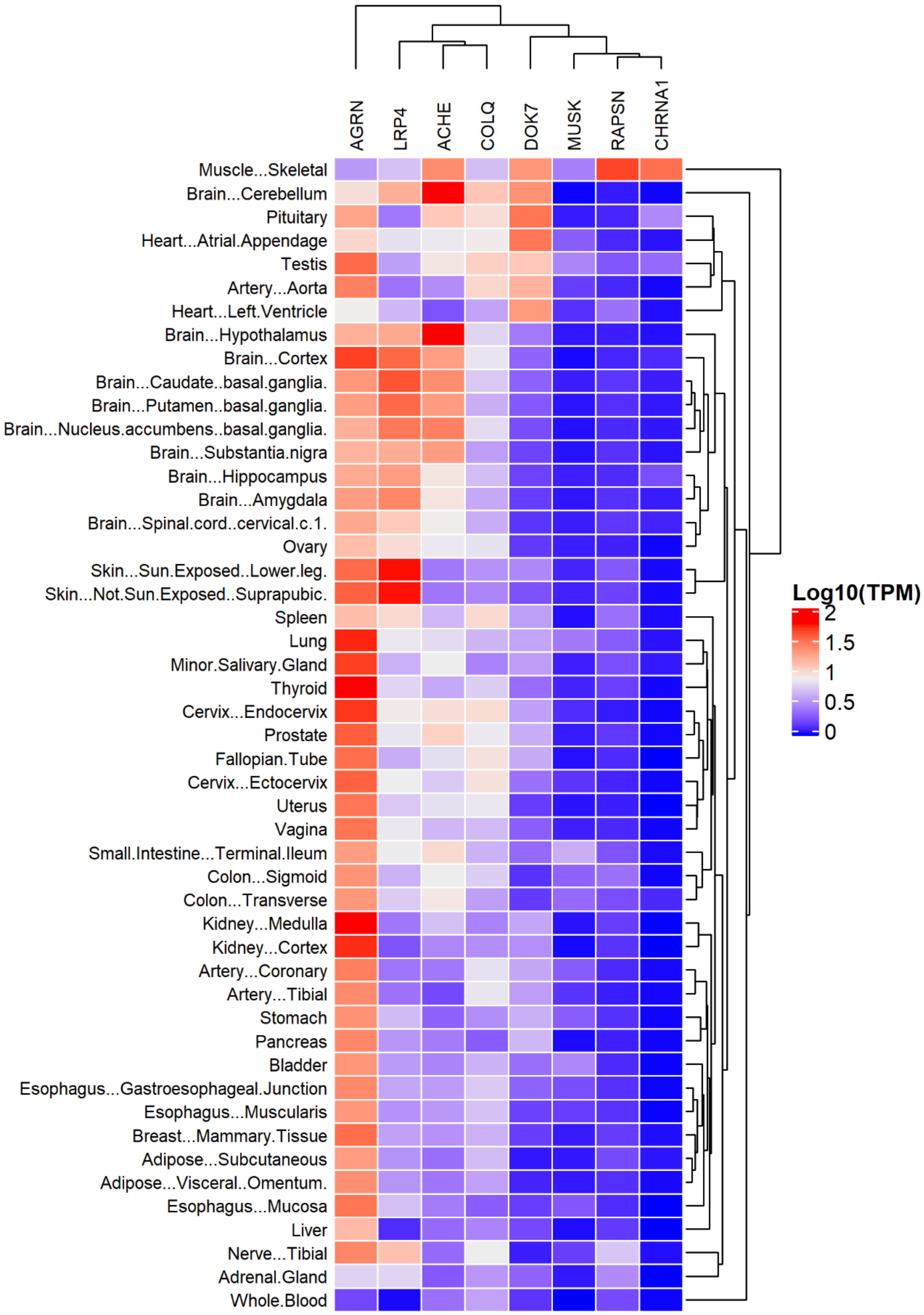


**Supplemental Fig. 2.** Hierarchical clustering of gene expression in the GTEx database using all isoforms of *ACHE* and *AGRN*.

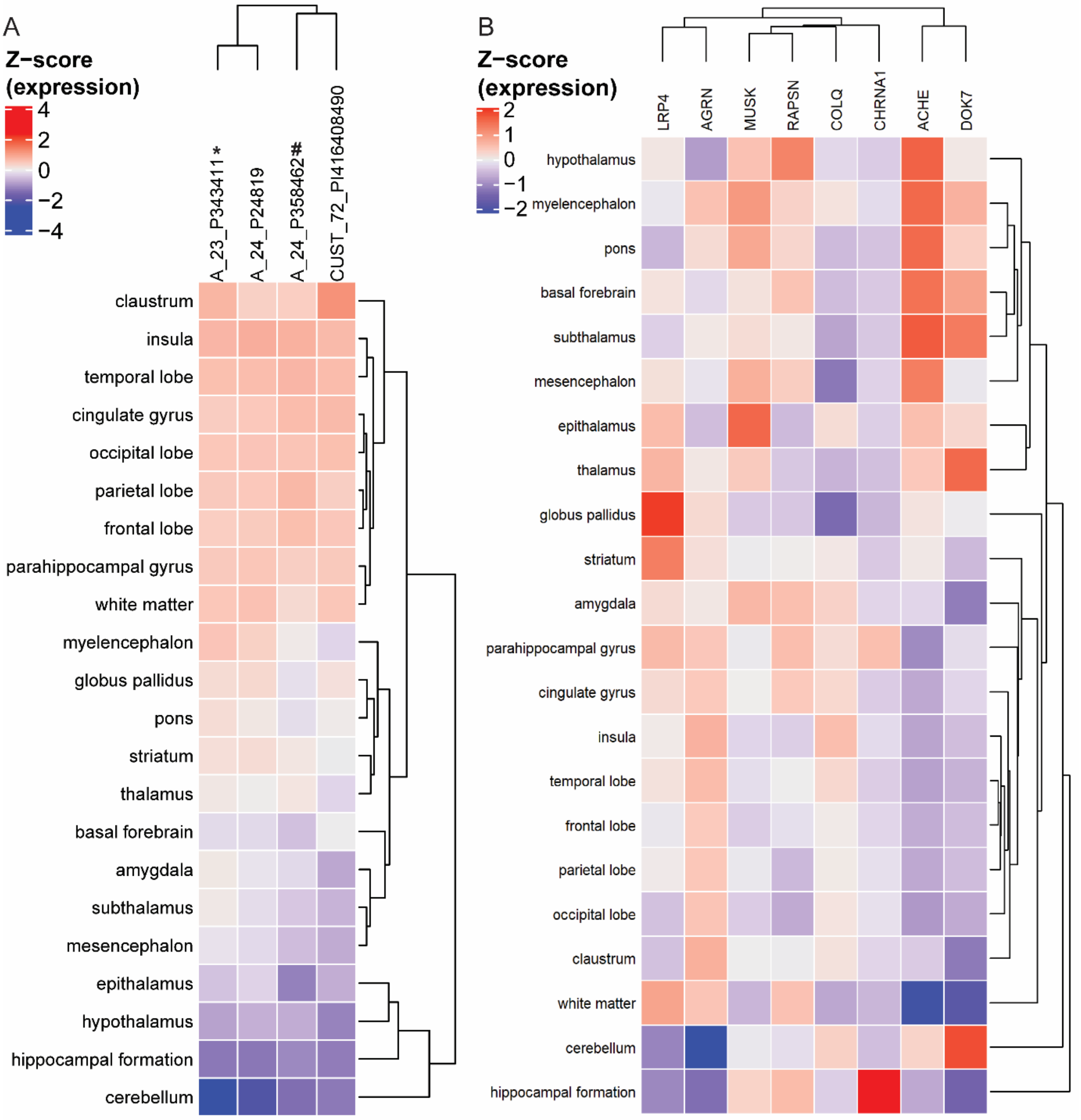


**Supplemental Fig. 3.** Heatmaps from Allen human brain atlas. Relative expression of different agrin probes across distinct brain areas (A). # probe that selectively detects NMJ-relevant isoform of agrin. * probe for AGRN used in the brainscope tool detecting all AGRN isoforms. Hierarchical clustering using probe for AGRN that does not discriminate between isoforms (*) (B).

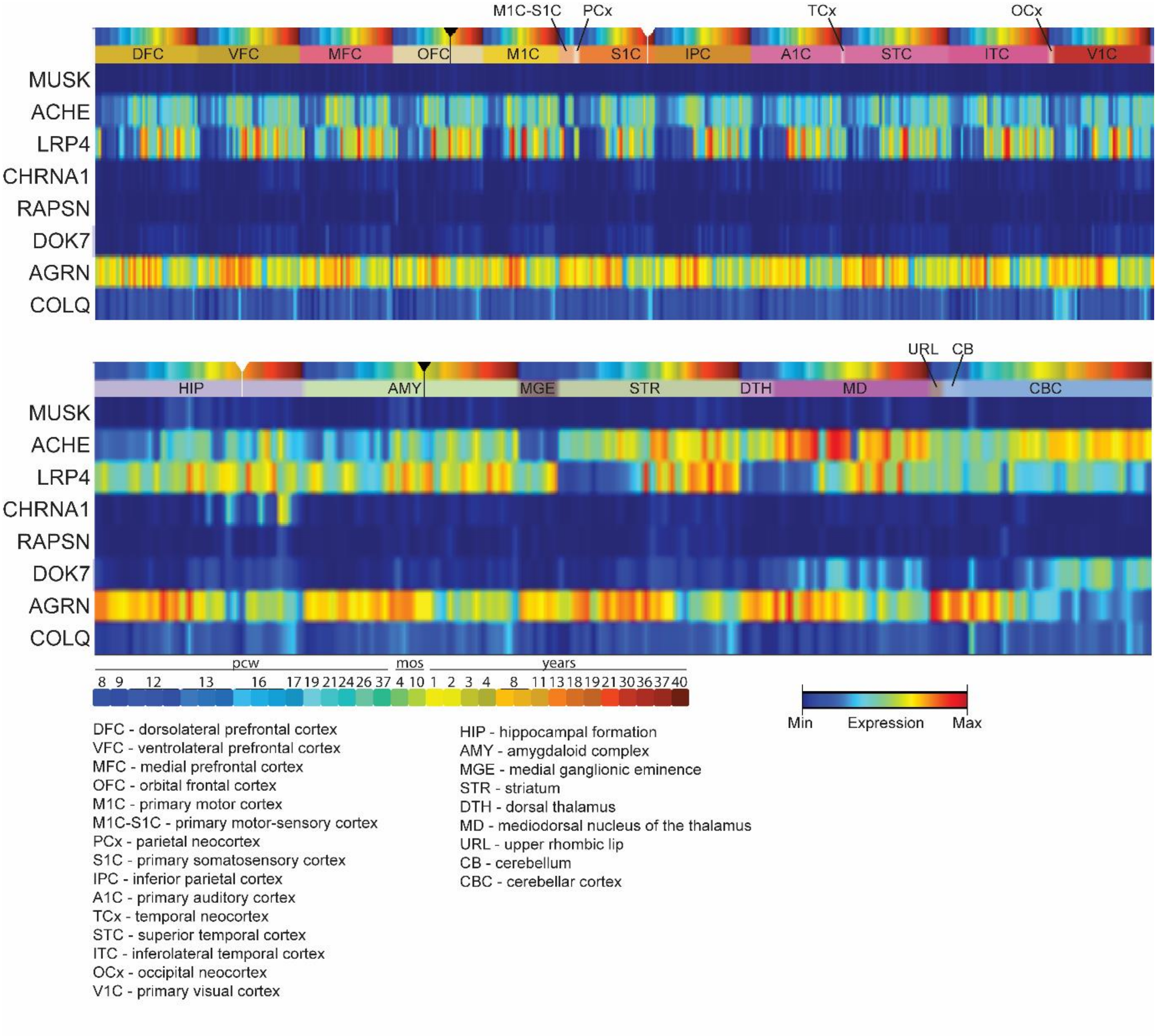


**Supplemental Fig. 4.** Gene expression of MG-related genes during human development do not show the same pattern across development. CHRNA1 expression is increased across development restricted to the hippocampus. *DOK7* shows a pattern of increasing expression across development, with highest expression in the cerebellum and thalamus. *MUSK* is lowly expressed during prenatal development in the hippocampus, amygdala and cerebellum. *RAPSN* shows very limited expression across development, restricted to some subcortical regions of the brain. Visualized using the BrainSpan portal.
